## Supplementary Information for "Cell swelling enhances canonical β-adrenergic signaling"

This document contains

- Supplementargy Figures 1-12
- Supplementary Table 1

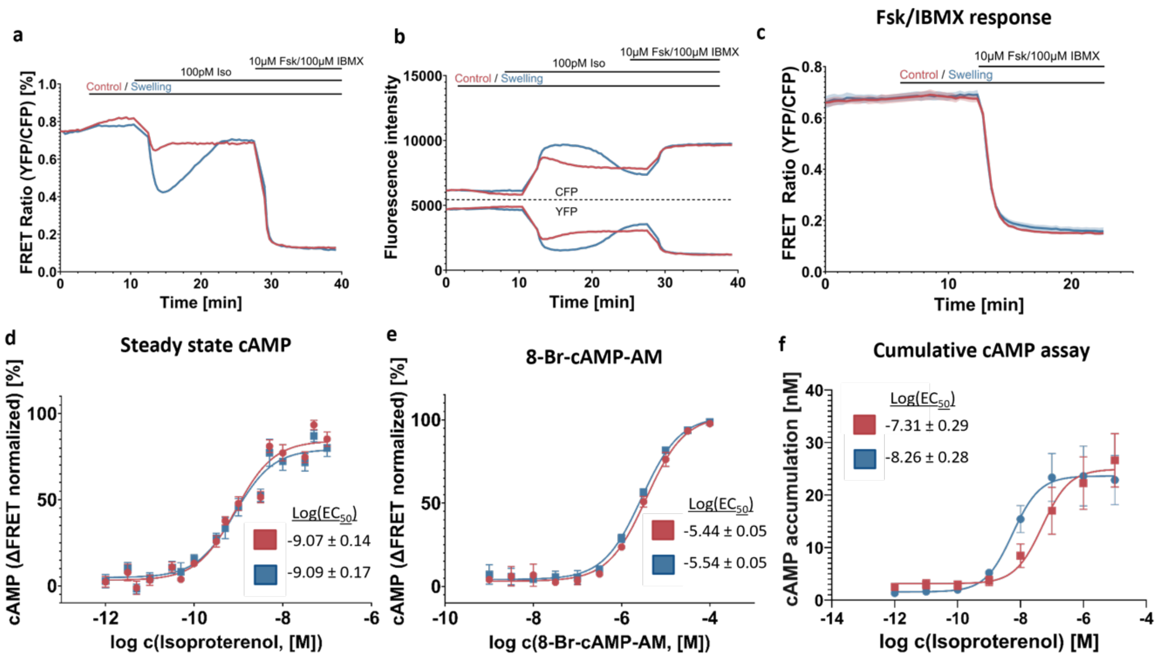

**Supplementary Fig. 1: Effect of swelling on receptor signaling is not caused by sensor artifacts.**

**a,b** non-normalized FRET ratio (**a**) and respective CFP and YFP channel traces (**b**) for the representative curve showed in the main **Fig. 1 c** (out of 8 independent experiments); **c** average normalized kinetic curves of acceptor/donor ratio measured at baseline, after medium change to control or swelling medium and after addition of 10 µM Forskolin + 100 µM IBMX (mean ± SEM; n = 3 independent experiments); **d** averaged concentration response curves representing steady state cAMP concentrations as indicated in **Fig. 1 c** (mean ± SEM; n = 7 independent experiments); **e** averaged concentration response curves of HEK293 Epac-SH187 stable cell line being stimulated by increasing concentrations of 8-Br-cAMP-AM (normalized to baseline and 10 µM forskolin + 100 µM IBMX; mean ± SEM; n = 3 independent experiments); **f** concentration-response curves obtained by measuring cAMP concentrations with the AlphaScreen endpoint cAMP assay in cell lysates of untransfected HEK293T cells, that were incubated in an isotonic and hypotonic medium containing indicated concentrations of isoproterenol for 10 minutes prior to cell lysis (mean ± SEM; n = 12 replicates from 4 independent experiments). Source data are provided as a Source Data file.

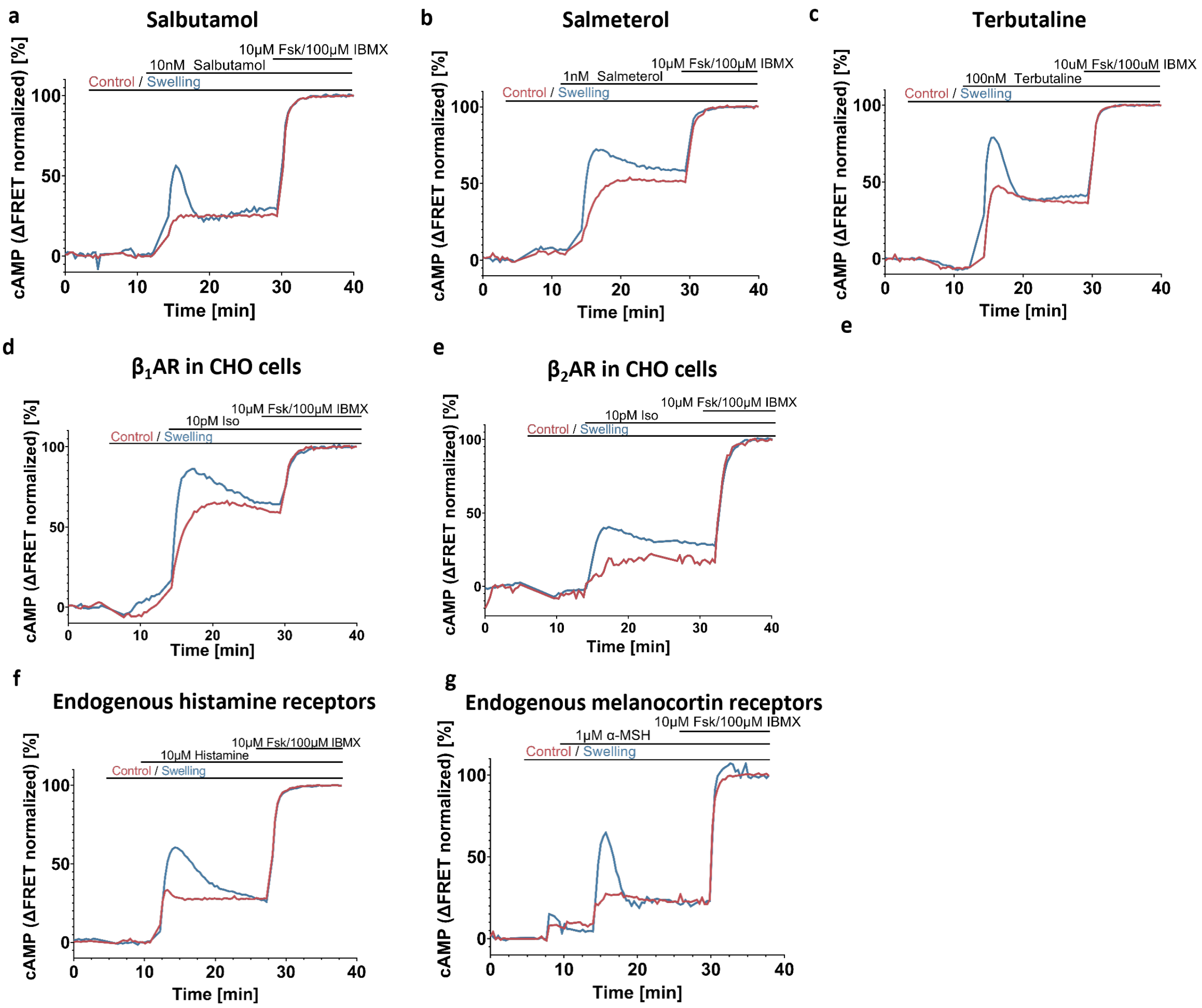

**Supplementary Fig. 2: Effect of swelling on receptor signaling is generalizable to other β-AR agonists, β-AR subtypes and to other Gs-coupled GPCRs.**

**a-c** representative curves (out of 3 independent experiments) showing kinetic cAMP concentrations as a response to different β-adrenergic agonists in HEK293 Epac-$S^{H187}$ stable cells (normalized to baseline and 10 µM Forskolin + 100 µM IBMX); **d,e** representative curves (out of 3 independent experiments) of intracellular cAMP concentration upon stimulation with 10 pM isoproterenol of CHO cells transfected with Snap-β1AR (or Snap-β2AR for **e**) and Epac-$S^{H187}$ sensor; **f,g** representative curves (out of 2 independent experiments) showing kinetics cAMP concentrations as a response to histamine (**f**) and α-MSH (**g**) in HEK293 Epac-SH187 stable cells (normalized to baseline and 10 µM forskolin + 100 µM IBMX). Source data are provided as a Source Data file.

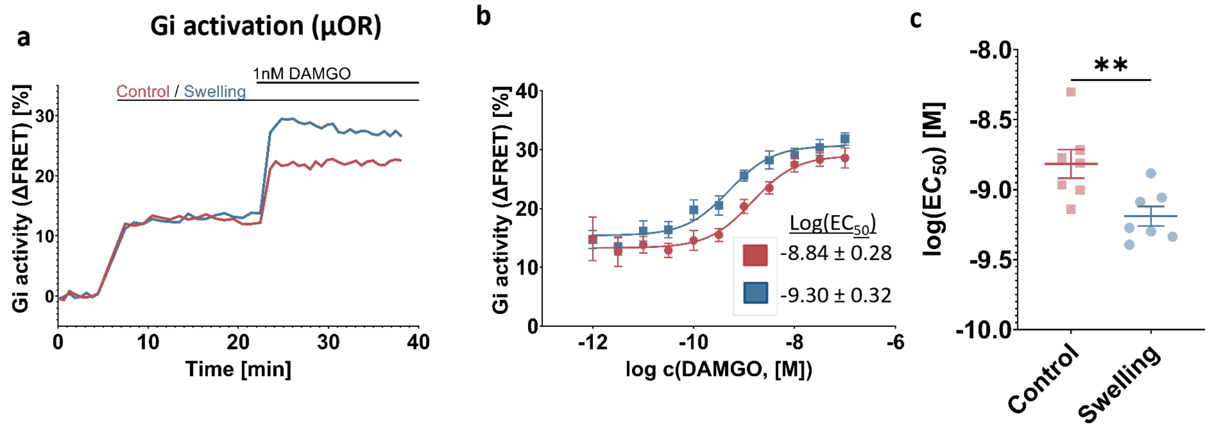

**Supplementary Fig. 3: Effect of swelling on receptor signaling is generalizable also to $G_i$-coupled receptors.**

**a-c** Representative curve (**a**), averaged concentration response curve (**b**) and log($EC_{50}$) values from individual experiments (**c**) comparing $G_{i2}$ FRET-sensor response to DAMGO in control and swollen HEK293 cells, transiently transfected with µOR and the $G_{i2}$-FRET sensor (n = 7 plates form 5 independent experiments; unpaired two-tailed t-test; p value = 0.0069). Source data are provided as a Source Data file.

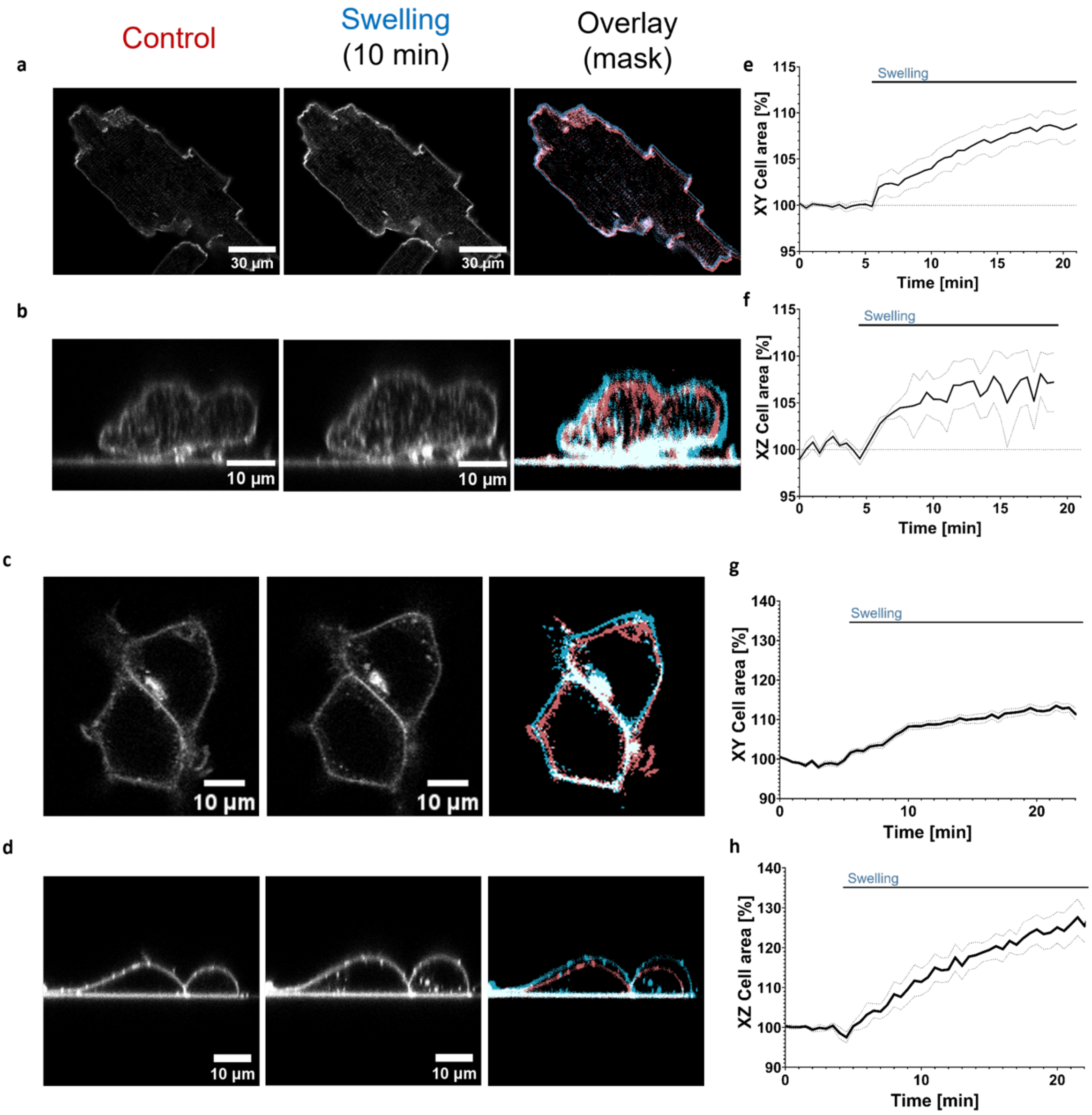

**Supplementary Fig. 4: Characterization of cardiomyocyte and HEK293 swelling**
**a-d** Representative confocal images of cardiomyocytes (**a,b**) and HEK293 cells (**c,d**) under isotonic conditions (left) and after 20 minutes of exposure to swelling medium (middle), with an overlay of cell edges under both conditions (right; isotonic in red and swelling in blue). Number of experiments conducted is reported below in the captions e-g. **a** and **c** show XY projections, while **b** and **d** show cells in a XZ projection; **e-h** average area changes over time for cardiomyocytes (**e,f**) and HEK293 cells (**g,h**) in the XY and XZ projections respectively (mean ±SEM, n = 6 cells from 3 independent experiments (**a**,**e**), n = 3 cells from 2 experiments (**b**,**f**), n = 35 cells from 3 experiments (**c**,**g**) and n = 7 cells from 3 experiments (**d**,**h**)). Source data are provided as a Source Data file.

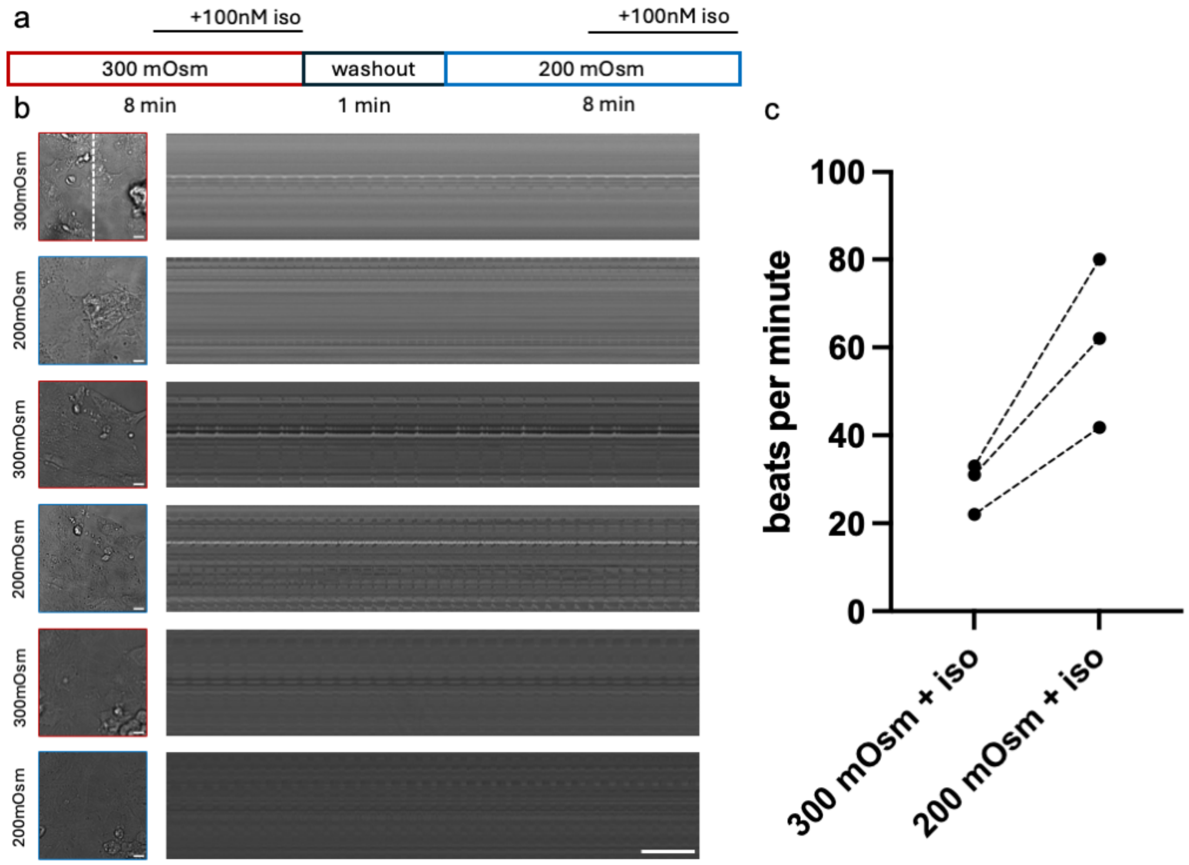

**Supplementary Fig. 5: Characterization of hiPSC-CM chronotropic response upon swelling**
**a** Schematic of the experimental setup. **b** Three wells of the same batch of hiPSC-CM were imaged in a brightfield microscope first in isosmotic (300 mOsm) and then hypotonic (200 mOsm) conditions upon addition of 100 nM iso. Kymograph were collected from line selections to determine the beating rate. **c.** Beating rate before and after swelling for the three sets of experiments. Source data are provided as a Source Data file.

Supplementary movie 1 (cAMP in CM upon Iso and Fsk/IBMX stimulation)

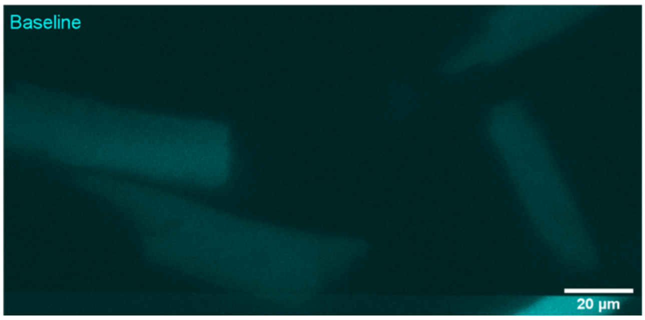

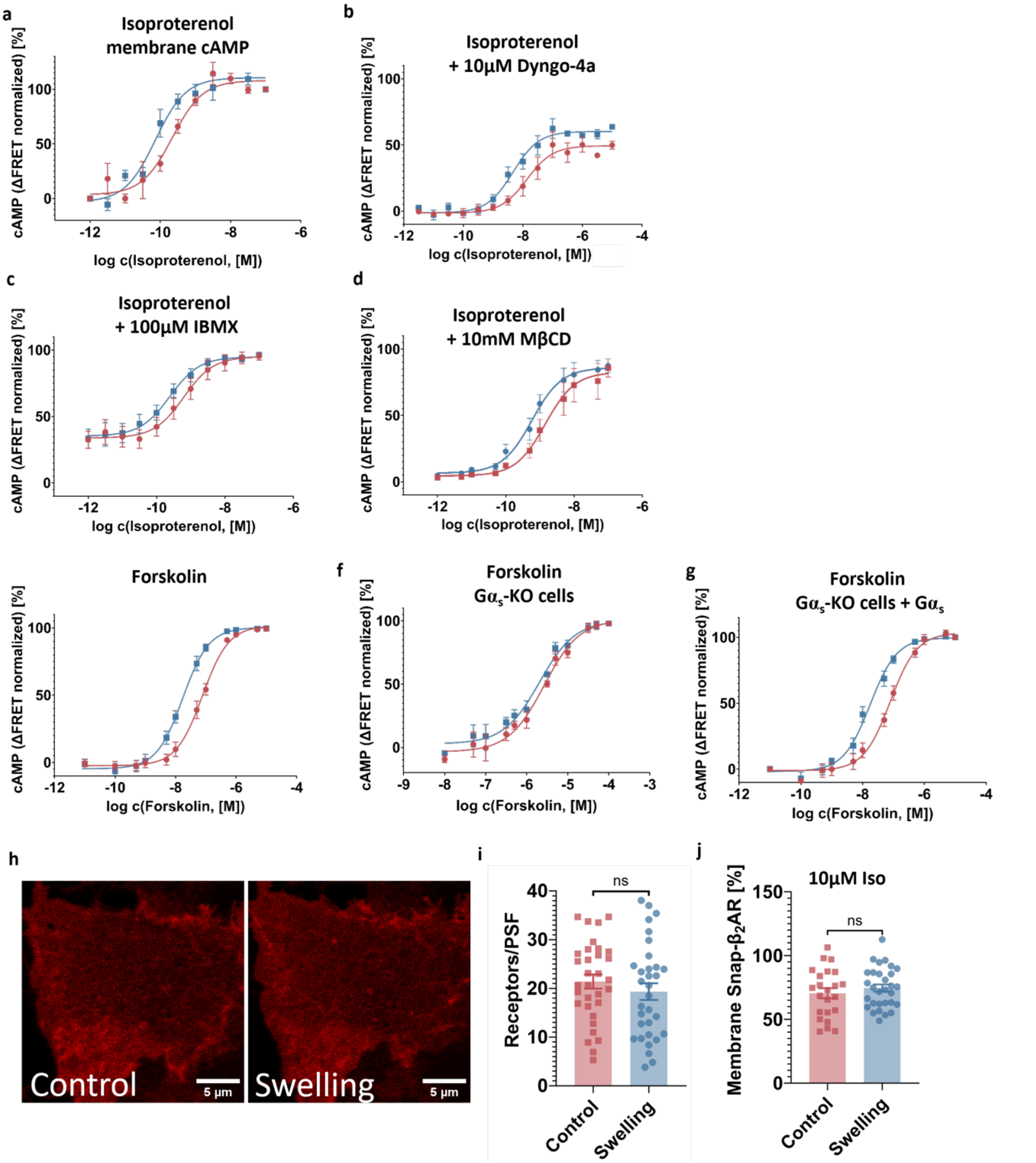

**Supplementary Fig. 6: Impact of cell swelling on isoproterenol potency under selective regulation of the β-adrenergic signalling cascade**

**a-g** Averaged concentration response curves of intracellular cAMP concentrations under following conditions (mean ± SEM; n = individual experiments): (**a**) local cAMP at the plasma membrane upon Iso stimulation measured with an Epac1-Camps-CAAX sensor (transiently transfected HEK293 cells; n=3); (**b**) cytosolic cAMP measured upon Iso stimulation after treatment with 10 µM Dyngo4a for 30 min (HEK293 Epac-$S^{H187}$ stable cell line; n= 4); (**c**) cytosolic cAMP measured upon Iso stimulation with simultaneous addition of 100µM IBMX (HEK293 Epac-$S^{H187}$ stable cell line; n=4); (**d**) cytosolic cAMP measured upon Iso stimulation after treatment with 10 mM methyl-β-cyclodextrine for 30 min (HEK293 Epac-$S^{H187}$ stable cell line; n=3); (**e,f,g**) cytosolic cAMP measured upon forskolin stimulation in (**e**) wt $G\alpha_s$ phenotype (HEK293 transfected with Epac-$S^{H187}$; n= 5), (**f**) $G\alpha_s$-KO cells (transfected with Epac-$S^{H187}$; n=4) and (**g**) $G\alpha_s$-KO cells transiently transfected with $G\alpha_s$ and Epac-$S^{H187}$ (n=5); **h** representative confocal images (number of experiments indicated below in the caption i-j) of basolateral membrane of HEK293AD cell transiently transfected with Snap-β2AR and labelled with SnapSurface-Alexa674 dye before and after hypotonic treatment; **i** number of receptors per pixel as obtained from single cells before and after hypotonic treatment (mean ± SEM; n = 32 cells from 4 independent experiments ;paired two-tailed t-test; p value = 0.1132); **j** relative number of receptors at the plasma membrane (compared to baseline) after 10 min of 10μM isoproterenol stimulation (mean ± SEM; n = 23 cells from 3 independent experiments; unpaired two-tailed t-test; p value = 0.3929). Source data are provided as a Source Data file.

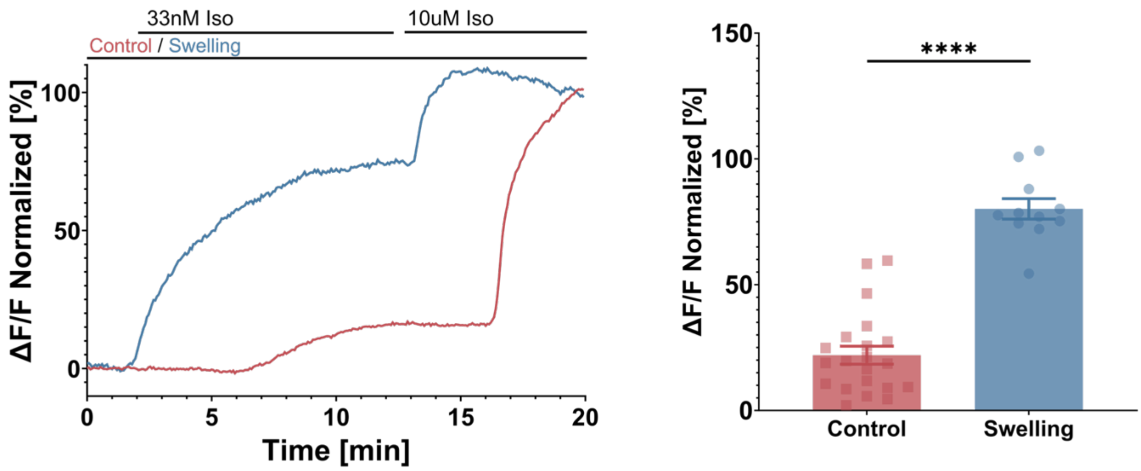

**Supplementary Fig. 7: Swelling leads to an increased β-arrestin2 recruitment independent of G$\alpha_s$**
**a** representative curve (out of 21 (control) and 11 (swelling) cells from 3 independent experiments) showing relative increase of membrane fluorescence collected as first 33 nM, and then at saturating 10 µM concentration of isoproterenol is added to the cell (normalized to baseline and 10 µM Iso response); **b** relative fluorescence increase measured upon β-arrestin2-eYFP recruitment upon 33 nM isoproterenol stimulation in swollen vs non-swollen cells (mean ± SEM; n = 21 (control) and 11 (swelling) cells from 3 independent experiments; unpaired two-tailed t-test; p value <0.0001). Source data are provided as a Source Data file.

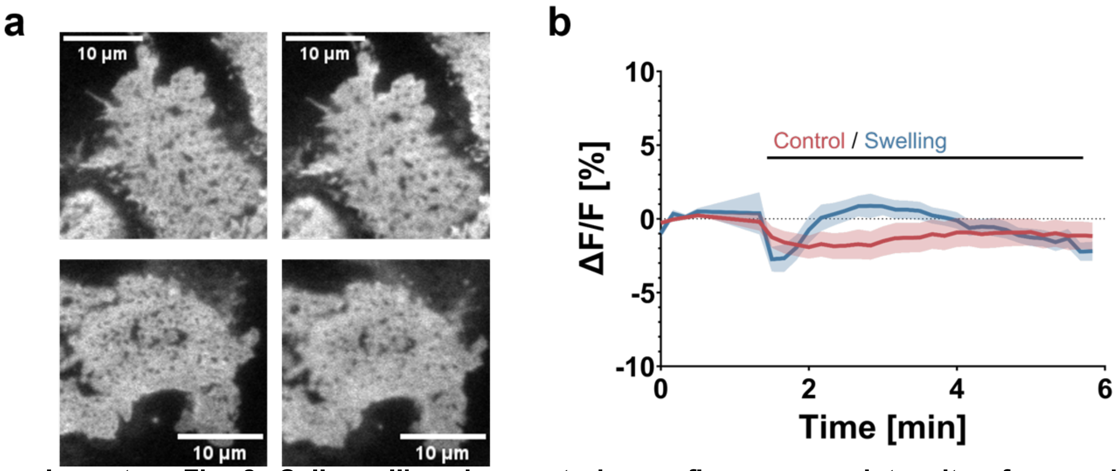

**Supplementary Fig. 8: Cell swelling does not change fluorescence intensity of a membrane anchored fluorescent protein under TIRF-M measurements.**
**a** Representative TIRF-M images (out of 11 cells from 2 independent experiments) of basolateral membrane of HEK293AD cell transiently transfected with Venus-CAAX before (left) and after (right) medium change for the control (upper) and swelling (lower) condition; **b** averaged curves of relative fluorescence intensity change upon medium replacement (mean ± SEM; n = 10 cells (control) and 11 cells (swelling) from 2 independent experiments). Source data are provided as a Source Data file.

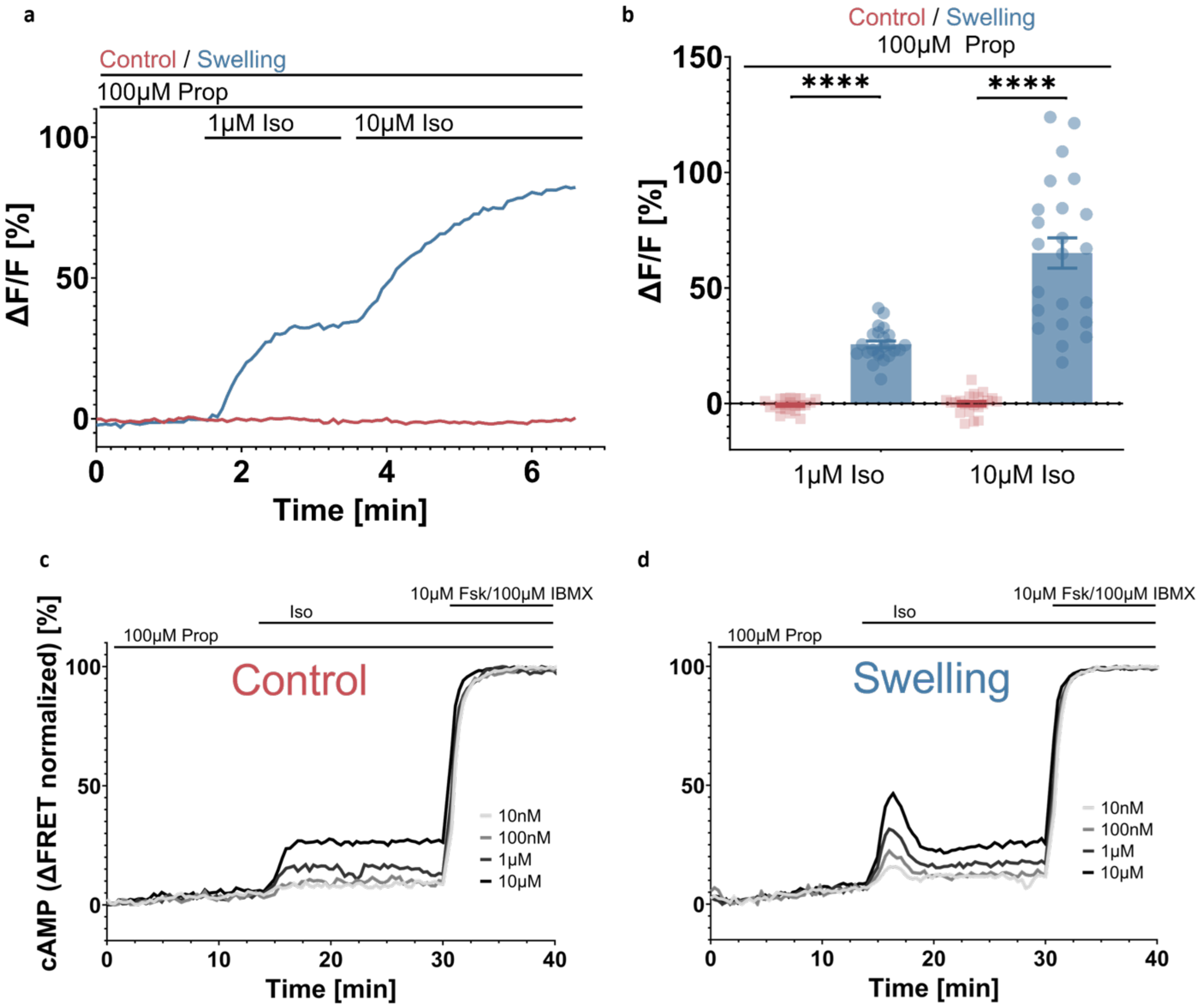

**Supplementary Fig. 9: Swelling reduces propranolol blocking efficiency of β2-AR**

**a** Representative time-sequence (out of 3 independent experiments) of the relative increase of Nb80-eYFP fluorescence signal at the basolateral membrane, as isoproterenol is added after 60 min incubation with 100 µM propranolol; **b** relative fluorescence increase measured upon Nb80-eYFP recruitment upon 1 µM and 10 µM isoproterenol stimulation in swollen vs non-swollen cells (mean ± SEM; n = 21 cells (control) and 23 cells (swelling) from 3 independent experiments; unpaired two-tailed t-test; p values: <0.0001 (1µM Iso), <0.0001 (10µM Iso)); **c, d** representative kinetic curves (out of 2 independent experiments) of acceptor/donor ratio measured in HEK293T cells stably expressing Epac-$S^{H187}$ in a microplate reader after 1h preincubation with 100 µM propranolol and consecutive addition of indicated isoproterenol concentrations; cells were additionally incubated in control (**c**) or swelling (**d**) medium, containing 100 µM propranolol, before isoproterenol addition. Source data are provided as a Source Data file.

Supplementary movie 2 (Nb37-eYFP recruitment)

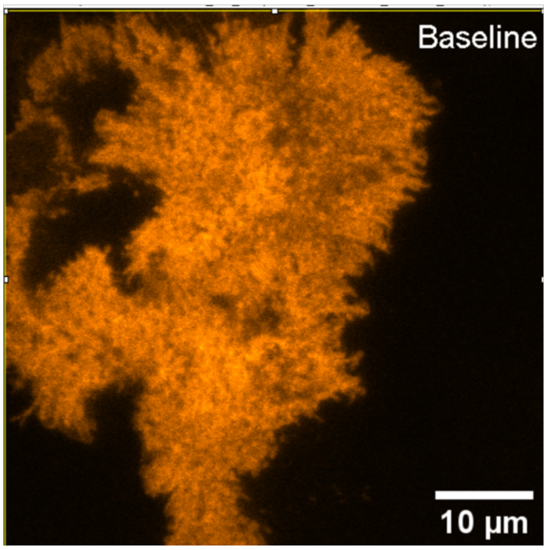

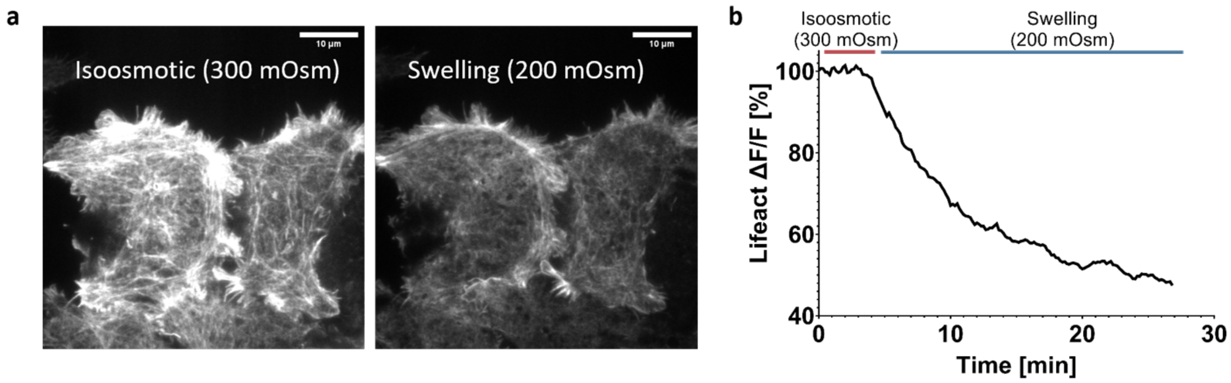

**Supplementary Fig. 10: F-actin labelling shows a decrease in cortical actin polymerization upon induction of cell swelling**

**a** Representative TIRF-M images (out of 3 independent experiments) of HEK293AD cells transfected with Lifeact-eGFP and visualizing F-actin in basal conditions (left) and after 20 minutes of incubation in swelling medium (right) **b** quantification of Lifeact-eGFP intensity over the course of time for one representative cell. Source data are provided as a Source Data file.

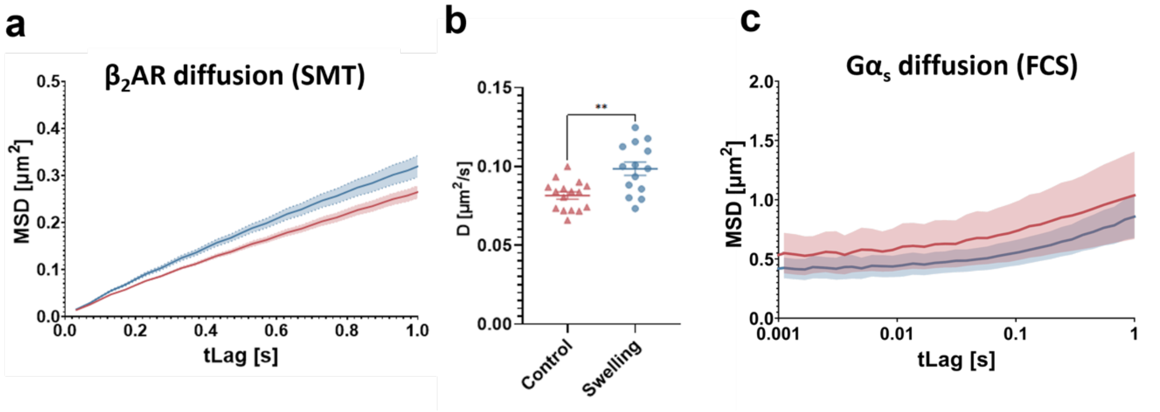

**Supplementary Fig. 11: Single molecule tracking shows faster ß2AR plasma membrane diffusion in swollen cells**

**a** averaged mean squared displacement curves obtained by tracking Snap-ß2AR diffusing on basolateral membrane of transiently transfected HEK293AD cells using TIRF-M; **b** diffusion coefficients calculated per cell from the first 0.15 s of corresponding MSD plots (MSD = 4Dt) (**a-b** mean ± SEM; n = 16 (control) and 14 (swelling) cells from 3 independent experiments; unpaired two-tailed t-test; p value = 0.0012); **c** averaged mean squared displacement curves obtained by tracking G$\alpha_s$-eYFP diffusing on basolateral membrane of transiently transfected HEK293AD cells using confocal microscopy-based linescan FCS (mean ± SEM; n = 9 cells from 2 independent experiments). Source data are provided as a Source Data file.

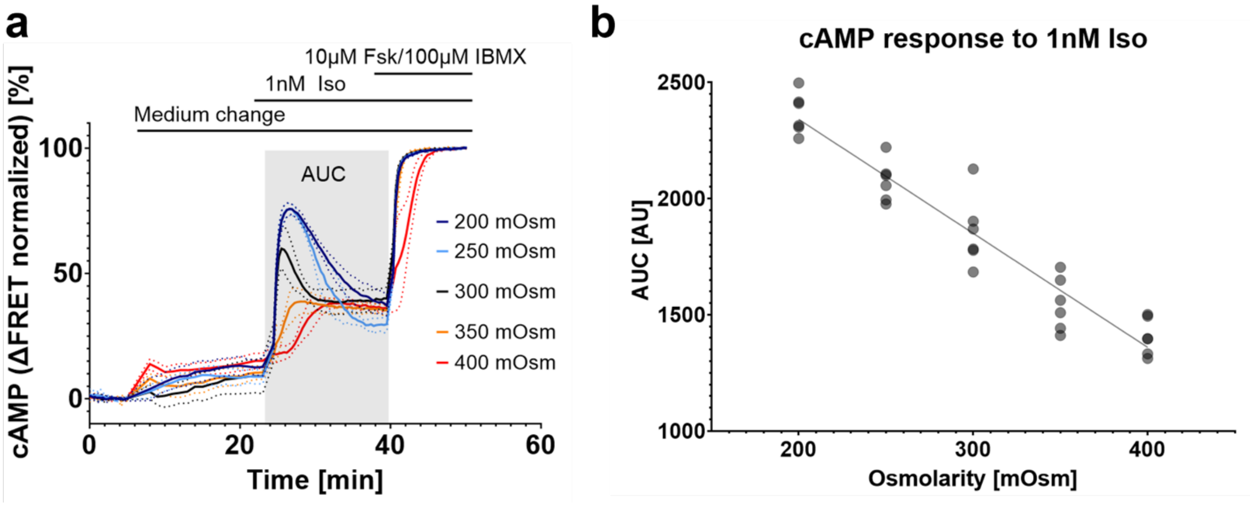

**Supplementary Fig. 12: Osmolarity can be titrated to differentially modulate cAMP production**
**a** averaged curves of intracellular cAMP concentration under different osmolarities (mean ± SEM; n = 6 plates from 2 independent experiments); **b** areas under the curve obtained from single experiments and plotted against osmolarity, including a linear regression fit (n = 6 plates from 2 independent experiments). Source data are provided as a Source Data file.

**Supplementary Table 1:** Primers used for molecular cloning of the Gβ$_1$-2A-Gγ$_2$-IRES-Gα$_s$ (tricistronic G$_s$) and Gβ$_1$-2A-Gγ$_2$-IRES-Gα$_s$-CFP (tricistronic G$_s$-CFP) plasmids

| Number | Sequence (5'-3') | Template DNA | Output |
|---|---|---|---|
| P1 | TTGATGAGTTTGGACAAACCACAACTAGAAT | pcDNA 3.1 | Intermediate |
| P2 | tgaaaaacacgatgataaGCGGCCGCTCGAGCATGC | pcDNA 3.1 | |
| P3 | ATTCTAGTTGTGGTTTGTCCAAACTCATCAA | pcDNA 3.1 | |
| P4 | ctggtcaagctcccccatAAGCTTGGGTCTCCCTAT | pcDNA 3.1 | |
| P5 | gcatgctcgagcggccgcTTATCATCGTGTTTTTCA | Gi$_3$-FRET sensor (van Unen et al. *PLOS One* 2016) | |
| P6 | aatcccggccctaccggtATGGCCAGCAACAACACC | Gi$_3$-FRET sensor | |
| P7 | atagggagacccaagcttATGGGGGAGCTTGACCAG | Gi$_3$-FRET sensor | |
| P8 | ggtgttgttgctggccatACCGGTAGGGCCGGGATT | Gi$_3$-FRET sensor | |
| P9 | tgaaaaacacgatgataaATGGGCTGCCTCGGCAAC | Gα$_s$-split CFP (Hynes et al. *Journal of Biological Chemistry* 2004) | Tricistronic G$_s$-CFP |
| P10 | gcatgctcgagcggccgcTTAGAGCAGCTCGTATTG | Gα$_s$-split CFP | |
| P11 | caatacgagctgctctaaGCGGCCGCTCGAGCATGC | intermediate | |
| P12 | TTGATGAGTTTGGACAAACCACAACTAGAAT | intermediate | |
| P13 | gttgccgaggcagcccatTTATCATCGTGTTTTTCA | intermediate | |
| P14 | ATTCTAGTTGTGGTTTGTCCAAACTCATCAA | intermediate | |
| P15 | tgaaaaacacgatgataaATGGGCTGCCTCGGGAAC | Gα$_s$-short (cDNA.org #GNA0SL0000) | Tricistronic G$_s$ |
| P16 | gcatgctcgagcggccgcTTATAGCAGCTCGTACTG | Gα$_s$-short | |
| P17 | cagtacgagctgctataaGCGGCCGCTCGAGCATGC | intermediate | |
| P18 | TTGATGAGTTTGGACAAACCACAACTAGAAT | intermediate | |
| P19 | gttcccgaggcagcccatTTATCATCGTGTTTTTCA | intermediate | |
| P20 | ATTCTAGTTGTGGTTTGTCCAAACTCATCAA | intermediate | |
